# Latent-TGF-β has a domain swapped architecture

**DOI:** 10.1101/2024.09.27.615443

**Authors:** Mingliang Jin, Robert Seed, Tiffany Shing, Yifan Cheng, Stephen L. Nishimura

**Affiliations:** Department of Biochemistry and Biophysics, University of California, San Francisco, San Francisco, USA; Department of Pathology, University of California San Francisco, San Francisco, CA, USA; Howard Hughes Medical Institute, University of California San Francisco, San Francisco, CA, USA

## Abstract

The multifunctional cytokine TGF-β is produced in a latent form (L-TGF-β) where a RGD containing homodimeric prodomain forms a “ring” encircling mature TGF-β shielding it from its receptors. Thus L-TGF-β must be activated to function, a process driven by dynamic allostery resulting from integrin binding the L-TGF-β RGD motif. Here we provide critical evidence that defines a domain-swapped architecture of L-TGF-β, an essential component in the dynamic allostery mechanism of L-TGF-β activation.

## Introduction

TGF-β is an essential multifunctional cytokine with diverse functions in morphogenesis, extracellular matrix (ECM) and immune homeostasis^1^. Understanding the regulation of TGF-β function is paramount to dissect the roles of TGF-β in disease and to facilitate targeting for therapeutic benefit. For TGF-β to function, it must be exposed or released from its latent complex to engage TGF-β receptors through a process known as activation that is mainly mediated by binding to several integrins^2,3^. Latent-TGF-β (L-TGF-β) is composed of a disulfide-linked homodimeric prodomain which contains integrin binding arm domains connected to a straitjacket domain with a lasso loop encircling the furin-cleaved mature disulfide-linked homodimeric TGF-β growth factor^4^ (Fig. 1a and b). The lasso loops are flanked by the prodomain α1 and α2 helices which form the remaining key components of the straitjacket that maintain latency (Fig. 1b)^5^. During biosynthesis, N-terminal cysteines on the loop preceding the α1-helices covalently link to cysteines in milieu molecules such as GARP or LTBP1 that have a binding interface stabilizing both the growth factor and the straitjacket^6^.

**Fig. 1.**
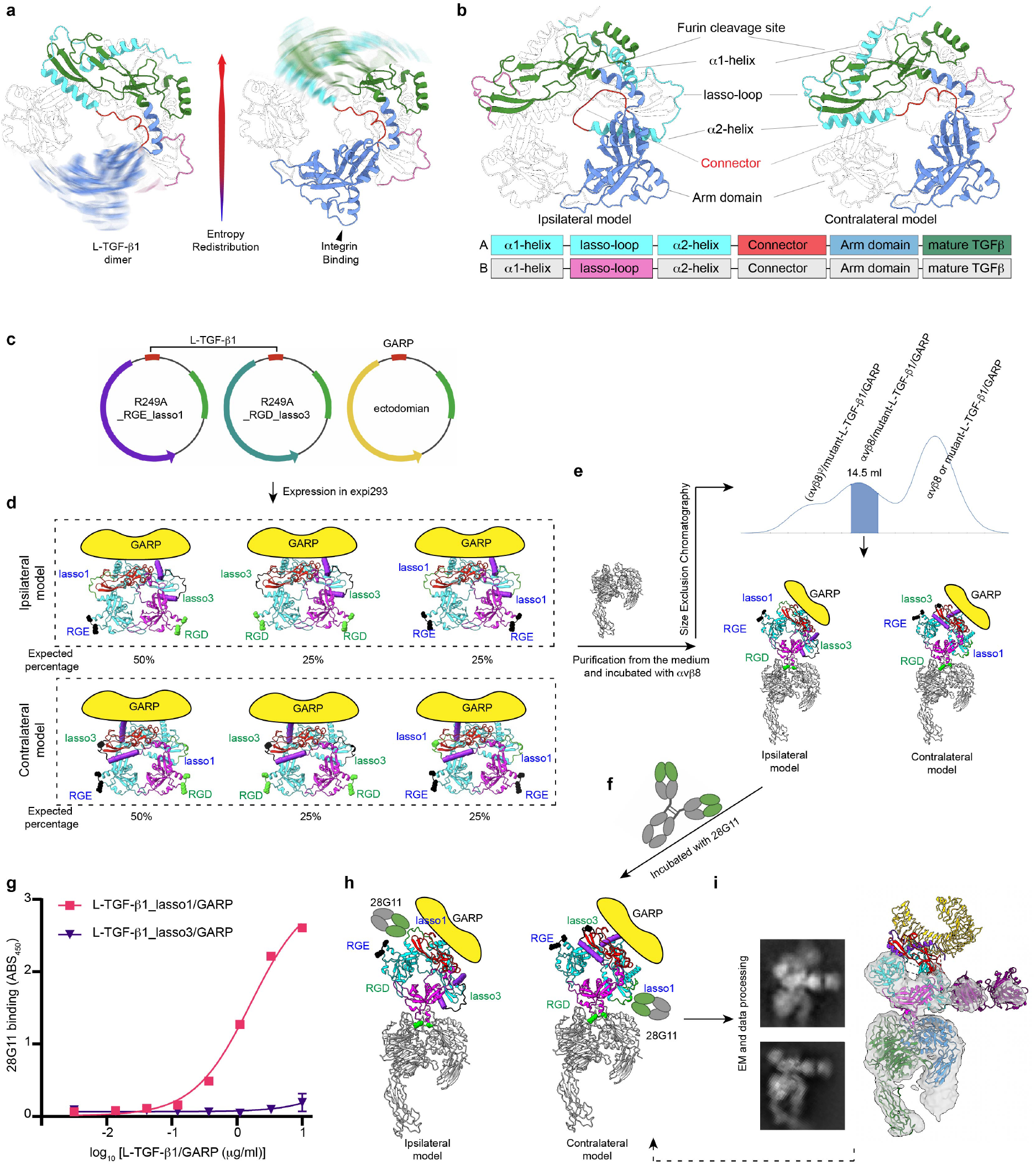
Experiment design and determination of domain architecture of L-TGF-β1/GARP. **a**, Ribbon diagram illustrating redistribution of conformational entropy in L-TGF-β1 from the RGD site to contralateral lasso loop prior (left) and after (right) αvβ8 binding. **b**, Ribbon diagram illustrating two possible architectures of L-TGF-β1 dimer, in which the straitjacket domain of each prodomain is connected ipsilaterally (left) or contralaterally (right) with the arm domain. Color scheme in ribbon diagram and domain arrangements are the same. **c**, Plasmid constructs of L-TGF-β1_R249A_RGE_lasso1 (left), L-TGF-β1_R249A_RGD_lasso3 (middle) and GARP (right). **d**, Anticipated expression products and their proportions from the 1:1:1 transfection of Expi293 cells. Ribbon diagrams of L-TGF-β1 in ipsilateral (upper) or contralateral (bottom) architectures are predicted by AlphaFold. **e**, SEC profile (middle) of purified mutant L-TGF-β1/GARP incubated with αvβ8 ectodomain (left). The shaded peak contains two possible complexes (bottom). **f**, Incubation of monoclonal antibody clone 28G11 with the SEC purified complex. **g**, ELISA confirms that 28G11 only binds lasso1 but not lasso3. Shown is a single experiment with error bars showing SD from 2 experimental replicates. **h**, Two models illustrating binding of 28G11 to mutant L-TGF-β1/GARP in ipsilateral (left) and contralateral (right) architectures. **i**, Representative 2D class averages and 3D reconstruction with atomic models of a generic Fab and contralateral L-TGF-β1/GARP docked.

We have recently shown that genetically engineering a loss-of-function mutation into this furin cleavage site results in mice that survive and are rescued from the lethal early tissue inflammation seen in *tgfb1* knock-out mice. These findings demonstrate that release of TGF-β from L-TGF-β is not required for its function^7^. For TGF-β activation to occur without release, TGF-β must be sufficiently exposed to interact with its receptors. Since the lasso loops of the straitjacket cover the receptor binding domains (RBD) on the tips of mature TGF-β, these loops must be sufficiently flexible after integrin binding to allow exposure of mature TGF-β. We found that binding of integrin αvβ8 to the RGD motif on the L-TGF-β arm domains induces flexibility preferentially to the contralateral straitjacket/lasso. This process is driven by dynamic allostery via redistribution of conformational entropy from the integrin binding site across the latent ring to increase the flexibility of the lasso loop (Fig. 1a). Increased flexibility of the lasso loop exposes the contralateral growth factor RBD to TGF-β receptors on the L-TGF-β/GARP expressing cell^7^. Defining the architecture of L-TGF-β is essential to understand how entropy is redistributed from the arm domain across the TGF-β latent ring to the contralateral lasso upon integrin αvβ8 binding. All available crystal structures of L-TGF-β assume the same architecture where the straitjacket/lasso loop is on the ipsilateral RGD-containing arm domain, but such architecture is assigned arbitrarily since the linker density was not resolved (Supplement Fig. 1a)^5,6,8,9,10^. AlphaFold predicts both non-swapped ipsilateral and swapped contralateral architectures (Fig. 1b). This ambiguity raises an essential question to understand the structural basis of entropy redistribution for L-TGF-β activation: Is entropy redistributed from the arm domain of one subunit to the lasso of the other subunit in an ipsilateral architecture, or within the same subunit in a domain swapped contralateral architecture? Here, we provide experimental evidence to definitely assign the domain swapped contralateral architecture to L-TGF-β.

## Results and discussion

In our recent study, we analyzed the cryo-EM structure of L-TGF-β1/GARP, where a weak density links the arm domain to the contralateral straitjacket domain (Supplement Fig. 1b, left)^7^, contradicting the presumed ipsilateral architecture (Fig. 1b, left). However, this density is insufficiently robust to definitively define the domain architecture. Thus, the domain architecture of L-TGF-β remains ambiguous due to the challenge of obtaining a high-resolution structure of this flexible loop region.

To define the domain architecture of L-TGF-β1, we created a system in which the architecture can be unambiguously identified without the need for a high-resolution structure. We generated two versions of an L-TGF-β1 expression plasmid (Fig. 1c): one with the intact RGD binding site and its TGF-β1 lasso loop (lasso1) replaced by the equivalent loop from L-TGF-β3 (lasso3), and the other with the intact lasso1 but containing a RGE mutation in its integrin binding motif. Both plasmids contain the R249A furin cleavage site mutation so that mature TGF-β1 remains covalently bound within the latent complex. We transfected these two plasmids in equal quantity together with GARP (Fig. 1c). The protein products, either in ipsilateral or contralateral architectures, consist of three versions of the L-TGF-β1 dimer, each covalently linked with a single GARP (Fig. 1d). One dimer is a mixture of two mutant monomers: one with a RGE motif and the other with a lasso3 loop. Thus, the RGD motif and lasso3 are either on the same side of the ring in the ipsilateral or on opposite sides in the contralateral architecture (Fig. 1d, left). The other two mutant dimers are symmetric (Fig. 1d, middle and right).

We incubated these purified L-TGF-β1/GARP mutants with αvβ8 followed by SEC to exclude mutant L-TGF-β1 with two RGE monomers that cannot bind integrin (Fig. 1e). We then added the antibody clone 28G11, which only recognizes the lasso1 loop^11^, to further identify the asymmetric dimer by single particle cryo-EM (Fig. 1f-i). The prediction was that 28G11 and αvβ8 bind this asymmetric L-TGF-β mutant on the opposite sides in the ipsilateral or the same side in the contralateral architecture (Fig. 1h). Indeed, 2D class averages and 3D reconstruction of the complex, even at a modest nominal resolution of 7 Å, clearly showed that 28G11 only binds to the lasso on the same side as the bound integrin. Docking the atomic models of L-TGF-β1/GARP and a generic Fab into this density map confirmed this assignment. Together, our experimental design with a clear logic provided a concise and straightforward result demonstrating the domain swapped contralateral architecture (Fig. 1i, and Supplement Fig. 1c). Thus, αvβ8 binding redistributes conformational entropy within the same monomer in the latent ring.

The domain-swapped architecture of L-TGF-β has mechanistic implications beyond αvβ8-mediated TGF-β activation. The integrin αvβ6 also binds to and activates L-TGF-β, but is hypothesized to involve actin-cytoskeletal force-induced deformation and release of mature TGF-β from L-TGF-β. This hypothesis was supported by molecular dynamics simulations based on models with an ipsilateral domain architecture^8^. These simulations, applying retrograde force through the β6 cytoplasmic domain, preferentially deformed the ipsilateral protomer, releasing mature TGF-β^8^. It will be interesting to see how dynamic simulations of force-induced αvβ6-mediated TGF-β activation are impacted by the domain-swapped architecture.

The domain-swapped architecture of L-TGF-β1 is likely to apply to other L-TGF-β isoforms, since the linker regions of L-TGF-β2 and L-TGF-β3 are of similar length to L-TGF-β1 (Supplement Fig. 1d). Indeed, we observe a weak density of the connector in L-TGF-β3/GARP^7^, consistent with the domain-swapped architecture (Supplement Fig. 1b, right). Interestingly, the TGF-β3 connector contains a cysteine that, based on its location, would easily form a disulfide linkage with the connector on the other monomer in the domain-swapped contralateral architecture, but likely remains unpaired in the non-domain-swapped ipsilateral configuration (Supplement Fig. 1d).

Among other TGF-β superfamily members, the domain-swapped architecture was clearly seen in the crystal structure of Activin A and was also assigned to myostatin, although a connector density was not observed (Supplement Fig. 1e)^12,13^. Our work shows an approach that may help in determining the domain architecture of myostatin and other TGF-β superfamily members, which all have prodomains that non-covalently link to their growth factors^14^. In conclusion, definitively defining the domain architecture of L-TGF-β and its related superfamily members is critical to understanding the mechanism of latency and activation.

## Methods

### Recombinant Protein Expression

L-TGF-β1_RGD_Lasso3 (where the A31-L44 in lasso1 loop was swapped with T31-V42 from the L-TGF-β3 lasso3 loop) has been previously described^7^. To produce L-TGF-β1_RGD_lasso3/L-TGF-β1_RGE_lasso1/GARP, Expi293 cells were transiently transfected with equal amounts of human L-TGF-β1_RGD_lasso3, L-TGF-β1_RGE_lasso1, and Strep-His-GARP plasmids as shown in Fig. 1c.

### Protein Production

The secreted ectodomain of αvβ8 integrin was produced by transfecting ExpiCHO cells with integrin constructs following the previous protocol^15^. After 5 days of growth, cells were centrifuged, and the supernatant was filtered through a 0.2 μm PES membrane (Millipore). Protein was purified from the supernatant via affinity chromatography using a Protein G column crosslinked with the 8B8 antibody, which binds to αv integrin^16^. Elution was achieved with 100 mM glycine (pH 2.5), followed by buffer adjustment and size exclusion chromatography (Superose 6 Increase 10/300 GL, GE Health-care) in 20 mM Tris-HCl pH 7.4, 150 mM NaCl, 1 mM CaCl_2_, and 1 mM MgCl_2_.

To produce secreted mutant L-TGF-β1/GARP, Expi293 cells were transiently transfected with three plasmids: L-TGF-β1_R249A_RGE_lasso1, L-TGF-β1_R249A_RGD_lasso3, and GARP ectodomain tagged with a Strep-His tag. The supernatant was collected by centrifuging the cell culture after 5 days of growth and then filtered through a 0.2 μm PES membrane. Protein purification was done using Ni-NTA agarose, followed by washing with a buffer containing 0.6 M NaCl, 0.01 M Tris (pH 8.0), and elution with 250 mM imidazole in TBS. The eluted protein was applied to a Strep-tactin agarose column and washed with TBS. To remove the tag, HRV-3C protease was added, and the mixture was incubated overnight at 4°C. Finally, the protein was concentrated to about 1 mg/ml in a TBS buffer using centrifugal concentrators.

Mutant L-TGF-β1/GARP and αvβ8 were first incubated at room temperature for 30 min, subjected to size exclusion chromatography, and correct peaks were pooled and concentrated to 0.31 mg/ml. 28G11 (Biolegend, San Diego, CA) was used without further purification.

### Cryo-EM

Purified mutant αvβ8/L-TGF-β1/GARP were incubated with 28G11 (1 mg/ml) at room temperature for 30 min at a molar ratio of 1:1. The final protein complex concentration was 0.37 mg/ml. For cryo-EM grid preparation, 3 μl of the complex was deposited onto Quantifoil 100 holey carbon films Au 300 mesh R 1.2/1/3. The grids were glow-discharged for 30 s at 15 mA prior to sample application and freezing. The complexes were frozen using a FEI Vitrobot Mark IV with a 1 s blot time. All grids were frozen with 100% humidity at 22 °C and plunge-frozen in liquid ethane cooled by liquid nitrogen.

**Supplementary Figure 1.**
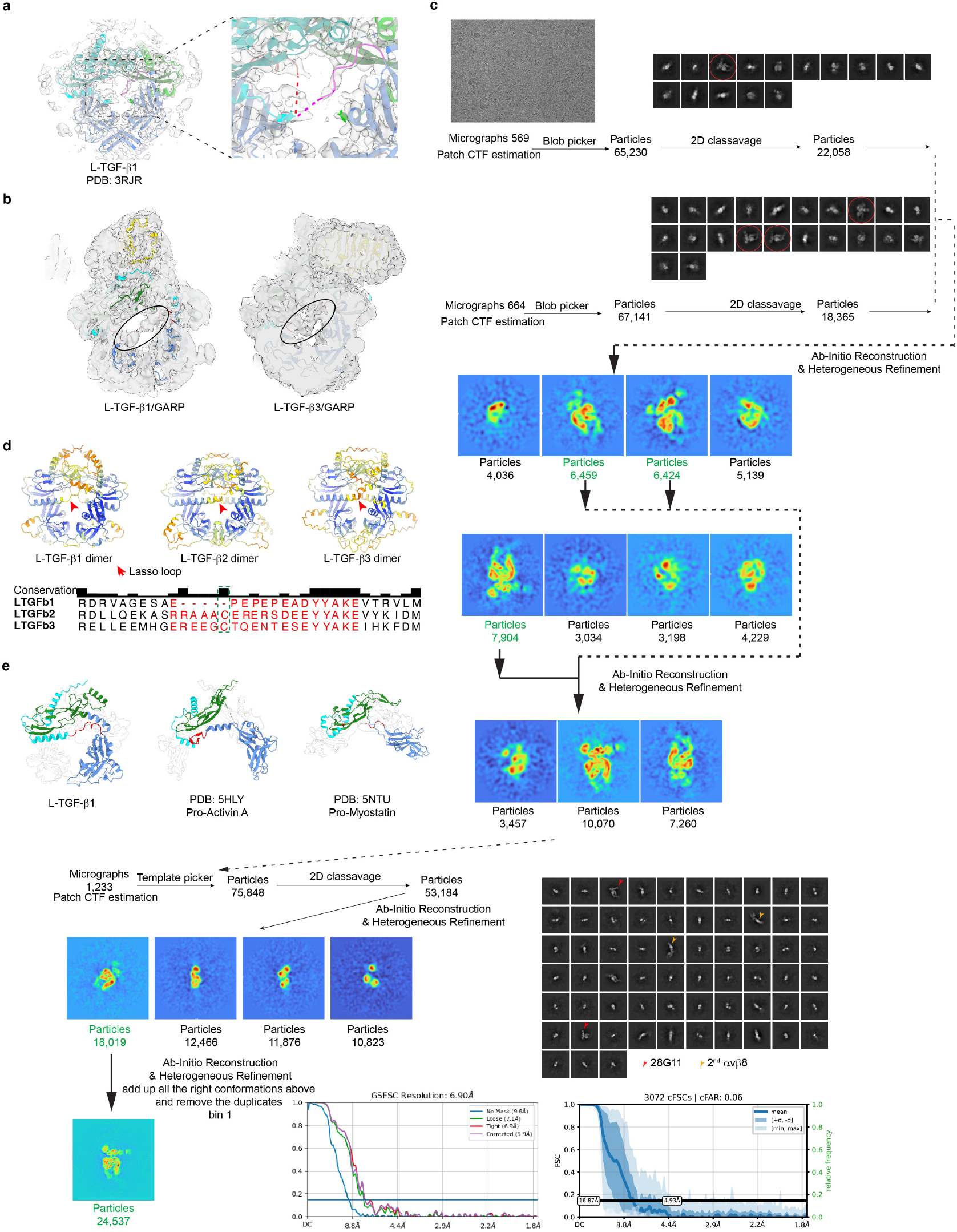
Single particle cryo-EM of mutant L-TGF-β1/GARP/28G11 complex. **a**, Calculated electron density map of L-TGF-β1 crystal structure (PDB: 3RJR) fitted with the atomic model. The enlarged view shows the density of the connector region is incompletely resolved, with the red dashed line linking the chains in an ipsilateral configuration. Magenta dashed line illustrates a possible alternative domain linkage for a contralateral configuration. **b**, Cryo-EM density map of L-TGF-β1/GARP (left) and L-TGF-β3/GARP (right) without sharpening^7^, shown at low contour level, showing a weak density (circled in black) suggesting the domain linkage for a contralateral configuration. **c**, Single particle cryo-EM data processing pipeline of αvβ8/L-TGF-β1/GARP/28G11. **d**, Alphafold prediction of L-TGF-β1, 2, and 3 in contralateral form, lasso loops are indicated, with the cysteines in TGF-β2 and -β3 in the green dashed box. **e**, Crystal structures of L-TGF-β1 (left), Pro-Activin A (PDB: 5HLY, middle), and Pro-Myostatin (PDB: 5NTU, right) highlighting the domain-swapped architecture with the connectors in red.

The data set was collected on a Thermo Fisher 200 KeV Glacios equipped with a GATAN K3 direct detector camera. 1,324 movies were collected at a nominal magnification of 69,000x, and the defocus range was set to be between -1.1 and -2.2 μm. The detector pixel size was 0.576 Å and the dose was 63 e^-^/Å^2^.

The data processing of αvβ8/L-TGF-β1/GARP/28G11 was carried out with CryoSPARC, with the workflow shown in Supplement Fig. 1c. The nominal resolution is estimated using the gold standard FSC = 0.143 criterion. Final reconstruction and directional FSC (cFSCs in CryoSPARC) show clear signs of anisotropic resolution, indicating that the dataset suffers from preferred orientation. Docking of the atomic model of αvβ8/L-TGF-β1/GARP and a generic Fab into the density map was performed using UCSF Chimera^17^. The location of 28G11 on L-TGF-β1 matches the previous cryo-EM structure of L-TGF-β1/GARP/28G11^11^.

### X-ray Map Density Calculation

The structure factor of L-TGF-β1 (PDB: 3RJR) was obtained from the PDB and converted to an mrc file which can be recognized by UCSF Chimera in COOT.

### AlphaFold Prediction

The predictions of human L-TGF-β1 dimers were performed using two identical TGF-β chains without signal peptide or templates by AlphaFold (https://colab.research.google.com/github/deepmind/alphafold/blob/main/notebooks/AlphaFold.ipynb)^18^.

### Sequence Alignments

Multiple protein sequence alignments for L-TGF-β were generated using Clustal Omega (https://academic.oup.com/nar/article/47/W1/W636/5446251).

### Antibody Binding Assay

ELISA plates were coated with serial dilutions of recombinant TGF-β1/GARP or recombinant TGF-β1_lasso3/GARP (10 μg/ml) in PBS for 1 hr at room temperature (RT). Wells were then washed in PBS and blocked with 5% BSA in PBS for 1 hour at RT. 28G11 or isotype control antibody was added (1 μg/ml) in PBS for 1 hr at RT. After washing in PBS with 0.05% Tween-20, bound antibodies were detected using anti-mouse-HRP and TMB substrate, with colorimetric detection performed using a Glomax Explorer (Promega).

## Acknowledgements

Equipment at the UCSF cryo-EM facility was supported by NIH grants S10OD020054, S10OD021741, and S10OD025881 and is managed by D. Bulkley and G. Gilbert. We thank M. Harrington and J. Li for computational support. This work was partially supported by NIH R01HL134183 and R01HL165175 (S.L.N. and Y.C.). Y.C. is an investigator of the Howard Hughes Medical Institute. BioRender was used for some figure preparations.

## Declaration of Interests

S.L.N. is on the scientific advisory board (SAB) of Corbus Pharmaceuticals, LLC. Y.C. is on the SABs of ShuiMu Bio-Sciences Ltd. and Pamplona Therapeutics.

## Data Availability

Cryo-EM map of mutant αvβ8/L-TGF-β/GARP/28G11 is deposited to EMDB with accession number EMD-47130.

## Notes

### Competing Interest Statement

S.L.N. is on the scientific advisory board (SAB) of Corbus Pharmaceuticals, LLC. Y.C. is on the SABs of ShuiMu BioSciences Ltd. and Pamplona Therapeutics.

